# Multitaper Estimates of Phase-Amplitude Coupling

**DOI:** 10.1101/2021.03.02.433586

**Authors:** Kyle Q. Lepage, Cavan N. Fleming, Mark Witcher, Sujith Vijayan

**Affiliations:** Virginia Tech, School of Neuroscience; Carilion Roanoke Medical Hospital

## Abstract

Phase-amplitude coupling (PAC) is the association of the amplitude of a high-frequency oscillation with the phase of a low-frequency oscillation. In neuroscience, this relationship provides a mechanism by which neural activity might be coordinated between distant regions. The dangers and pitfalls of assessing phase-amplitude coupling with existing statistical measures have been well-documented. The limitations of these measures include: (i) response to non-oscillatory, high-frequency, broad-band activity, (ii) response to high-frequency components of the low-frequency oscillation, (iii) adhoc selection of analysis frequency-intervals, and (iv) reliance upon data shuffling to assess statistical significance. In this work, a multitaper phase-amplitude coupling estimator is proposed that addresses issues (i)-(iv) above. Specifically, issue (i) is addressed by replacing the analytic signal envelope estimator computed using the Hilbert transform with a multitaper estimator that down-weights non-sinusoidal activity using a classical, multitaper super-resolution technique. Issue (ii) is addressed by replacing coherence between the low-frequency and high-frequency components in a standard PAC estimator with multitaper partial coherence, while issue (iii) is addressed with a physical argument regarding meaningful neural oscillation. Finally, asymptotic statistical assessment of the multitaper estimator is introduced to address issue (iv).

## 1 Introduction

The coordination of oscillations of differing frequencies in neural circuits is thought to mediate inter-regional neural communication and facilitate learning through neural plasticity. One such form of between-frequency coordination is known as phase-amplitude coupling (PAC). Phase-amplitude coupling is a relationship between the phase of a low frequency oscillation and the amplitude of a higher frequency oscillation. It is implicated in learning and memory [1–5], in pathology [6–13] and in neuromodulation therapies [14] and is important in cognition [15–21]. For instance, both coupling between hippocampal sharpwaves and ripples and coupling between neocortical slow waves and sleep spindles are both thought to be important for sleep-mediated memory consolidation [22, 25–28]. Fig. (1) depicts an example of cross-frequency coupling between slow waves and sleep spindles in an intracranial recording.

**Figure 1:**
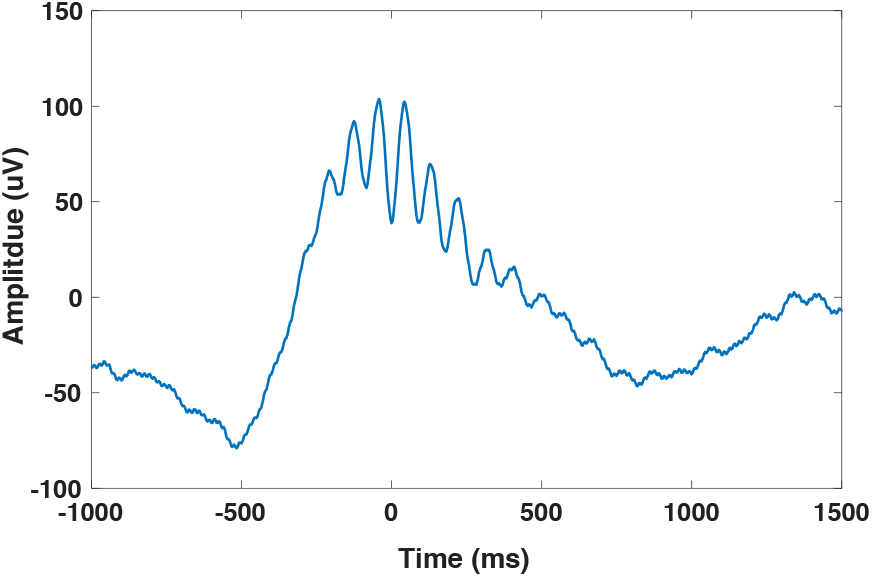
An example of phase-amplitude coupling (PAC). The occurrence of high-frequency oscillatory activity (11 Hz - 16 Hz) occurring in an intracranial recording taken from an epileptic patient during stage 3 Non-REM sleep is used to compute a time-locked average (depicted). An occurrence of this high-frequency activity during sleep is known as a sleep spindle. They are most prevalent during the upward deflection of the lower frequency oscillations (also shown). The exact spindle timing with respect to lower frequency oscillations probably depends on their frequency and location (see [22]). Sleep-spindles are important in sleep science: their prevalence is correlated with sleep-enhanced memory consolidation [23, 24].

Despite the potential importance of cross-frequency coupling (CFC) in understanding neural circuits, a number of issues related to the statistical assessment of phase-amplitude coupling using existing estimators have been reported [29–38]. These issues have lead to a flurry of activity described in [33, 39–52]. The statistical properties of four of these diverse methods have been assessed in [53]. Phase-amplitude coupling is extended to the transient setting in [54], and to the case of multivariate recordings in [55].

In this work, a novel, nonparametric phase-amplitude coupling estimator is introduced. This estimator is developed from ideas associated with the classical multitaper method of spectrum estimation. The proposed estimator is robust to non-sinusoidal, high-frequency activity, and further, it allows for the separate assessment of two types of cross-frequency coupling; namely the separation of (i) cross-frequency coupling where oscillation timing is stereotyped (i.e. phase-phase coupling (PPC)), from (ii) cross-frequency coupling where the envelope of the higher-frequency oscillation is stereotypically timed to the phase of the lower-frequency oscillation (i.e. phase-amplitude coupling, (PAC)). Thus, the case of non-sinusoidal, low-frequency oscillations that contain contaminating high-frequency components in their Fourier representation is handled. This mitigates the concerns raised in [35, 36]. Further, robustness to high-frequency but non-sinusoidal activity means that the proposed phase-amplitude coupling estimator is robust to broadband impulses occurring at a consistent phase of the low-frequency oscillation.

The multitaper method uses the optimal in-band energy concentrated discrete prolate spheroidal sequences^1^ (DPSSs) as data tapers applied prior to estimating Fourier-domain related quantities. This tapering confers improved sampling properties [57, 58] to nonparametric estimates of quantities involved in spectral analysis. These properties transfer to the proposed PAC estimator (see Section (7.2)). An issue common to all existing PAC estimators is the selection of frequency intervals to analyze. For the proposed estimator, interval selection is motivated based upon physical considerations and is specified as the number of cycles for which the high-frequency oscillation amplitude is expected to remain constant. Finally, reliance on data shuffling techniques to assess statistical significance is relaxed through the use of an asymptotic distribution (see Section (6)).

This paper begins with reviews of the multitaper method, and of phase-amplitude coupling, in Sections (2.1) and (2.2). The proposed multitaper PAC estimator is introduced in Section (3). In Section (7.2), the performance of the proposed estimator is demonstrated upon simulated data. Also using simulated data, in Section (7.1), robustness to contaminating impulses is demonstrated upon key quantities related to the proposed PAC estimator, while in Section (7.3), separation of phase-phase coupling from phase-amplitude coupling is demonstrated for non-sinusoidal pulse trains phase-locked to the amplitude of higher-frequency oscillation, but not to the phase of higher-frequency oscillation, which is random. In Section (8) the proposed PAC estimator is applied to real, intracranial data obtained from epileptic patients and compared against a classical PAC estimator that uses the Hilbert transform. The paper concludes with a discussion in Section (9).

## 2 Background

### 2.1 The Multitaper Method of Spectrum Estimation

The multitaper method of spectrum estimation was introduced in a sequence of papers beginning with [57] and including [59–66] and has been applied and further developed in the context of neuroscience [67–71]. The foundation of this method is the multitaper spectrum estimate. This estimate is computed as the average of direct-type nonparametric spectrum estimates, each of which is computed using a different discrete prolate spheroidal sequence taper [57]. The resulting spectrum estimate is characterized by (i) a reduction of harmful bias and, (ii) reduced variability due to the implicit averaging across small frequency intervals associated with tapering.^2^ While the foundation, the multitaper spectrum estimate is only part of the method. In particular, the multitaper method makes a minimum number of assumptions regarding the nature of recorded data, employs robust estimates to detect the presence of data features that contradict assumed properties, and then flags spurious data for further analysis with tailored methods. In Section (3), a multitaper estimator of phase-amplitude coupling is introduced. It is applied to simulated data in Section (7), and to actual intracranial data in Section (8).

### 2.2 Phase-Amplitude Coupling

Phase-amplitude coupling exists between two oscillations when the phase of the lower-frequency oscillation can be used to predict the phase of the instantaneous amplitude of a higher-frequency oscillation. Fig. (2) shows an example of phase-amplitude coupling. The instantaneous amplitude of the high-frequency oscillation, referred to as *γ* in Fig. (2), increases and decreases in-phase with the low-frequency oscillation (*θ* in Fig. (2)). PAC differs from phase-phase coupling (PPC), depicted in Fig. (3), in that the phase of the low-frequency oscillation, e.g. *θ* in Fig. (3), can be used to predict the phase of the high-frequency oscillation (*γ* in Fig. (3)). It is possible for two oscillations to exhibit both PPC and PAC simultaneously. This scenario occurs if a non-sinusoidal low-frequency oscillation couples with a higher-frequency oscillation oscillating at a frequency associated with non-zero coefficients in the Fourier representation of the non-sinusoidal, low-frequency oscillation. This is discussed further in Section (5), where multitaper across-frequency partial coherence is used to estimate PAC involving such a non-sinudoidal low-frequency oscillation.

**Figure 2:**
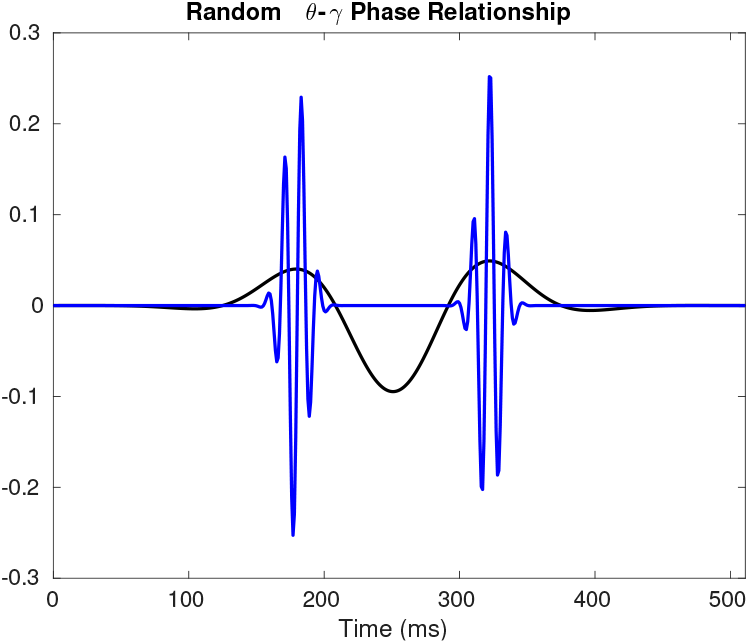
Simulated phase-amplitude coupling (PAC) between *γ* and *θ* rhythms.

**Figure 3:**
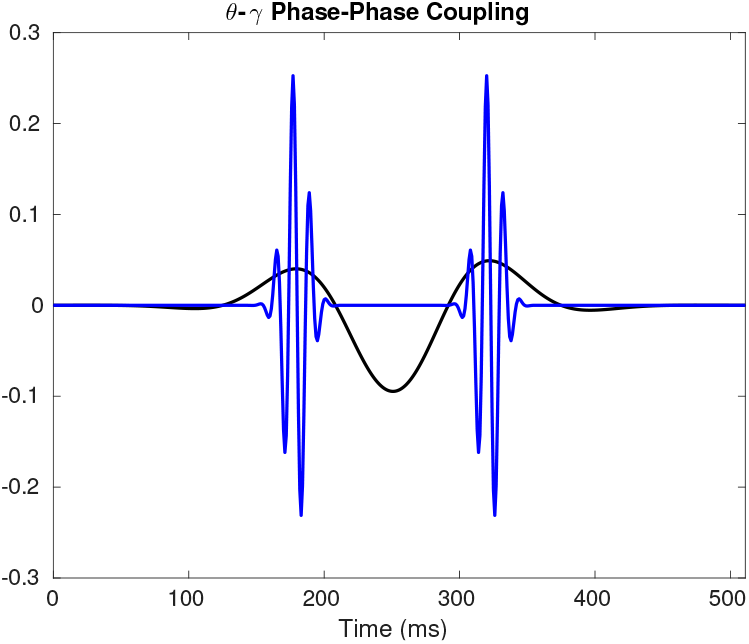
Simulated phase-phase coupling (PPC) between *γ* and *θ* rhythms. In this example, both phase-phase coupling and phase-amplitude coupling (PAC) are exhibited. This scenario, where both PPC and PAC exists, can be spurious and due solely to a non-sinusoidal, low-frequency, oscillation that consists of frequency components that exceed the frequency of the non-sinusoidal oscillation.

Estimation of the PAC, 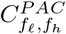, between frequencies *f*_*l*_ and *f*_*h*_ from the time-series *x* involves the following steps:

1. Select the bandwidth, *B*_*l*_, of the low-frequency component.
2. Bandpass filter *x* to the frequency interval (*f*_*l*_− *B*_*l*,_ *f*_*h*_ + *B*_*l*,_) to obtain *x*^(*l*)^
3. Select the bandwidth, *B*_*h*_, of the high-frequency component.
4. Bandpass filter *x* to the frequency interval (*f*_*h*_ *– B*_*h*,_ *f*_*h*_ + *B*_*h*_) to obtain *x*^(*h*)^
5. Estimate the instantaneous amplitude of *x*^(*h*)^. Typically, for time-index *t*, the instantaneous amplitude, 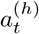, is computed as

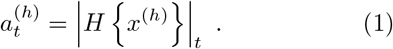 Here, *H* {*x*}_*t*_ is equal to the Hilbert transform of *x* evaluated at time-index *t*.
6. Assess the correlation between the phase, *ϕ*^(*l*)^, and the amplitude, *a*^(*h*)^. Various methods of doing this exist. One method calculates coherence. Another method computes an estimate of *ϕ*^(*l*)^ and then estimates and assesses the distribution of *a*^(*h*)^ conditioned upon this phase estmate.

Recent results have focused upon improving Step (vi) above conditioned upon estimates of *ϕ*^(*l*)^, and *a*^(*h*)^. The methods used include coherence, mutual information, and generalized linear modeling. A parametric approach that differs from these methods uses a generalization of autoregressive time-series modeling [45]. In this work, multitaper coherence is used to assess PAC between *x* and a multitaper estimate of the instantaneous amplitude (see Section (4)). A natural question pertains to the selection of the bandwidths, *B*_*f*_ and *B*_*h*_. In this work, the selection of *B*_*h*_ is based upon physical characteristics, allowing the remaining unknown bandwidth parameter, *B*_*f*_, to be specified through exploratory analysis, as is typical in classical spectral analysis.

## 3 Multitaper Estimate of Phase-Amplitude Coupling

Let *y* be a set of neural recordings. For trial *tr* and time-index *t*, let the corresponding measurement equal *y*_*t,tr*_. Without loss of generality set the sample period, Δ, to equal one second. Let *n* be the number of samples in each of the *N*_*tr*_ trials comprising *y*, the set of neural recordings. As an example, *y* might correspond to the recordings from a single EEG electrode across *N*_*tr*_ trials of an experiment. Let *v*^(*k*)^ be the *k*^*th*^-order discrete-prolate spheroidal sequence (DPSS). Following [57], the *j*^*th*^ eigencoefficient for the *tr*^*th*^ trial, evaluated at frequency *f*, is equal to,

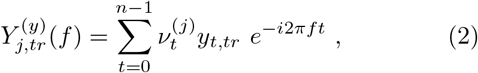

and a multitaper estimate *Š* of the spectrum is equal to,

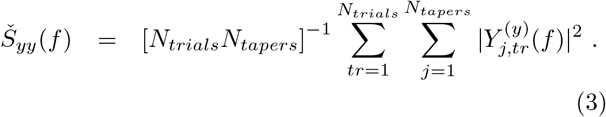

Given the multitaper estimate 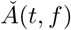 of the instantaneous amplitude of *y* evaluated at frequency *f* and time-index *t*, a multitaper estimate 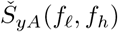 of the cross-spectrum between the frequency *f*_*l*_ of *y* and the frequency *f*_*h*_ of the instantaneous amplitude of *y* is equal to,

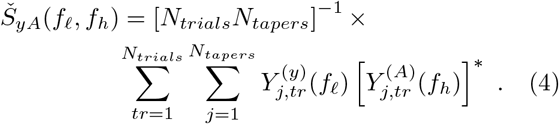

Here, *Y* ^(*A*)^ are the eigencoefficients computed by replacing *y* in Eqn. (2) with 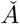 This multitaper estimate of the instantaneous amplitude *A* is introduced in Section (4). The proposed multitaper estimate of PAC, 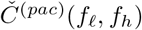, is equal to,

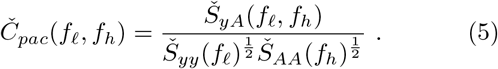

The performance of the proposed estimator is demonstrated in simulation in Section (7.2) and on intracranial recordings in Section (8). In a multiple trial setting, or in a single-trial setting using the Welch analysis method where sections of data are analyzed individually and then averaged, coherence is preferred to the phase-locking value; due to the down-weighting of noisy phase estimates in the computation of a coherence estimate [72].

### 3.1 PAC Phase Estimate: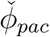

A multitaper estimate of the PAC phase is equal to,

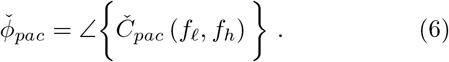

## 4 Multitaper Estimate of Instantaneous Amplitude, 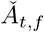

In [74, 75] estimates of the instantaneous amplitude of narrow-band filtered time-series are introduced that use the optimal DPSS sequences to improve accuracy. Here, we use such an estimate of the instantaneous amplitude, but modify it to down-weight the estimate if data are non-sinusodal. This latter weighting is computed from the P-values of the hypothesis test for harmonic modes (i.e. sinusoidal oscillations) first presented in [76].

The derivation begins by introducing time-dependence into the *k*^*th*^ eigencoefficient, *Y*_*k*_, defined in Eqn. (2) to obtain the *k*^th^ time-dependent eigencoefficient, 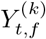, evaluated at time-index *t* and frequency *f* :

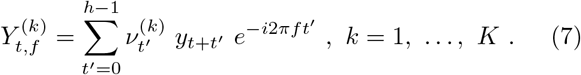

Here, *K* is the number of DPSS tapers, and *h* is the number of samples in the local interval: [*t, t* + *h* − 1]. Let the time-dependent amplitude, frequency and phase of a sinusoidal oscillation be equal to *A*_*t*_, *f*_*t*_, *ϕ*_*t*_, respectively. Substituting this model into Eqn. (7), results in a relationship between the amplitude *a*_*t*_ and the eigencoefficients (see A). The multitaper estimate, 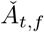, specified in Eqn. (29) is the least-squares estimate of *A*_*t*_ in this relation.

### 4.1 Physical Parameter Specification

Multitaper spectral estimates smear signal energy across intervals of length 2*W* and do not produce clearly distinguishable peaks that are spaced less than 2*W* apart in frequency. Consequently, the resolution in frequency associated with 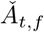 decreases with increasing DPSS parameter *W* It is desirable to specify a small, positive valued *W* parameter. On the other hand, the temporal resolution of 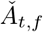 decreases with decreasing *W* for a fixed time-bandwidth parameter (*NW*), while the number of useable tapers increases with increasing *NW*. For example, setting *NW* to 4 results in about 6 tapers with signal energy predominantly within the frequency interval (−*W, W*). There exists a trade-off between frequency and temporal resolution; it results from the well-known signal processing time-frequency uncertainty relation. Physiologically interesting high-frequency phenomena are constant over a number of oscillation cycles. This duration is the temporal scale over which neural phenomena changes and guides our selection of temporal resolution and hence of *W*. Specifically, in this work the number of constant cycles is set to equal 3, and *h* is set equal to 3 oscillation periods, i.e. *h* = 3*/f*. Thus, for *NW* equal to 4, *W* = 4*f* /3, and the resulting frequency resolution of 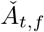 is inversely proportional to *f*. This inverse *f* dependence is consistent with the insight reported in [30]: to follow fast changes in instantaneous amplitude, a larger bandwidth is required. Further, this specification is consistent with *B*_*h*_ exceeding *f*_*l*_ for *f* > 3*f*_*l*_/4 as recommended in [34]. The number of tapers is chosen to be those with greater than or equal to ninety-nine percent of their signal energy within the analysis frequency interval (−*W, W*). See [77] for an interesting, if slightly tangential, discussion on time-frequency signal concentration.

### 4.2 Resistance to Narrowband, Non-sinusoidal Contamination

Spurious energy within a frequency interval can result from non-sinusoidal modulation in PAC, or from a temporally-punctate, and hence broadband, pulse. In either case, significant phase-amplitude coupling may result yet fail to indicate the expected relationship between oscillations. Resistance to such activity is a desirable feature for a PAC estimate. Such resistance can be accomplished by down-weighting 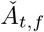 if non-sinusoidal activity is present. To accomplish this, the signal-energy explained within the Hilbert space spanned by the set of DPSSs can be compared to the signal-energy that is not explained. This procedure is detailed in B, where the frequency-dependent weight, wf is introduced in Eqn. (37). By estimating this weight for each interval of time, [t, t + h-1], at the desired frequency, f, time and frequency dependence is introduced. This weight appears in the proposed multitaper instantaneous amplitude estimate, Eqn. (29), where its scaling serves to reduce the amplitude estimated from narrowband, non-sinusoidal activity The resistance of 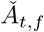 to non-sinusoidal activity is demonstrated in Section (7.1).

### 4.3 A Priori Bandpass Filtering

The estimate 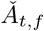 is capable of following a signal comprised of a single oscillation. If more than one oscillation is present then (i) the estimated instantaneous amplitude is not meaningful, and (ii), the computed weight, *w*, (see Eqn. (37)), is a value near zero. To avoid this, as with most PAC estimates, a priori bandpass filtering is required. In this work, the passbands are specified according to the frequency intervals associated with the named neural rhythms, i.e., delta, theta, alpha, beta, low-gamma, high-gamma, etc.

## 5 Multitaper Analysis of Non-Sinusoidal PAC

An important consideration when analyzing phase-amplitude coupling is whether or not the coupling is due to self-coupling arising from a non-sinusoidal oscillation. Such an oscillation is periodic with some period, *T*, and frequency *T* ^−1^, but does not have a sinusoidal shape. This implies that the waveform cannot be formed by simply repeatedly copying one period of a sinusoid and shifting it by one extra period to form the next period of the oscillation. Examples in neuroscience include the triangle-shaped theta rhythm often observed in hippocampal LFP recordings, and the sawtooth wave, commonly observed in electroencephalography recordings taken from sleeping participants just prior to or during a REM episode. To represent these oscillations, it is required that a number of sinusoidal oscillations are present with a specific phase relationship. This is an example of cross-frequency coupling – specifically, of phase-phase coupling. This phase-phase coupling provides the basis for a mitigating correction to the multitaper PAC estimate. Since phase-phase coupling is a linear relation between sinusoids in the sinusoidal (that is, Fourier) representation of the waveform, it suffices to decompose high frequency eigencoefficients into a sum of two components, a first component that is the phase-phase coupled component, and a second component that is not phase-coupled to the low frequency, modulating sinusoidal oscillation. Let the *j*^*th*^ element of the frequency-dependent vector of eigencoefficients **Y**_*f*_ equal 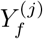. Then, for *f*_*l*_ < *f*,

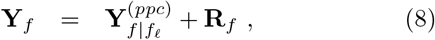

where 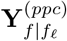 is the component predictable from **Y**_*fl*_ and **R**_*f*_ is the component that is not predictable. Therefore, **R**_*f*_ contains only phase-amplitude coupling with *f*_*l*_ that is not explained by phase-phase coupling, and thus, cannot be explained by a non-sinusoidal oscillation. The proposed correction to the multitaper PAC estimate, Eqn. (5), is to estimate **R**_*f*_ in Eqn. (8), and replace the eigen coefficients, 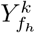, *k* = 1, …,*K* used in the computation of Eqn. (5) with the elements of the estimate **Ř**_*f*_ of **R**_*f*_ .This estimate is computed as the plug-in linear minimum-mean square error estimate of **R**_*f*_ :

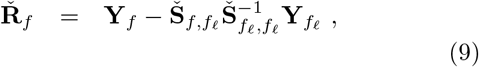

where the cross spectral matrix estimate is equal to,

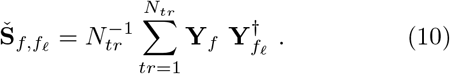

Similarly, the spectral matrix estimate is equal to,

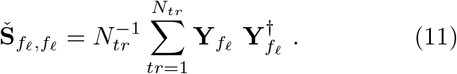

### 5.1 Intermittent PAC Modulated Pulses

Assessment of phase-amplitude coupling in the situation where modulating and modulated pulses are intermittent requires special consideration. This is the scenario, for example, with respect to slow waves and sleep spindles during slow-wave sleep [28, 78]. This situation is simulated in a few ways in Section (7). Let *x* be such an intermittent pulse, and let the *M* random pulse occurrence times be *τ*_*m*_, *m* = 1, …, *M* .Let *d* be the recording of this sequence of pulses:

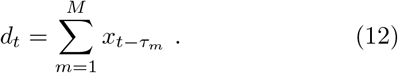

The *k*^*th*^ eigencoefficient computed from *d* is equal to,

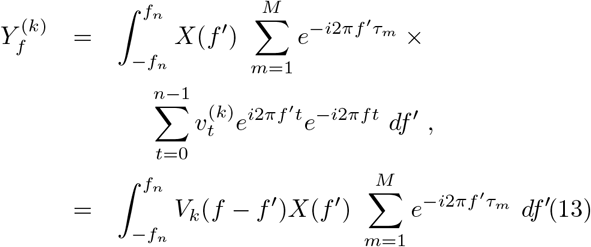

The random factor 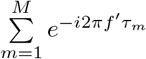 is real-valued in same-frequency coherence estimates, since they involve conjugate products that cancel the non-unity phase, leaving only the magnitude-squared of the sum. Thus, this factor does not affect same-frequency coherence. However, in across-frequency products, the factor *G*(*f*_*l*_, *f*_*h*_), is introduced,

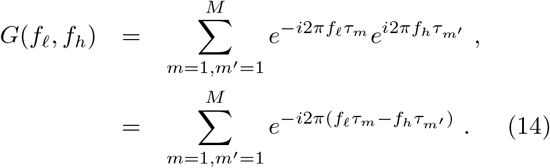

This factor is approximately equal to *M* in magnitude; however, the phase is random on the interval [*π, π*) radians and adds noise to the the calculation of the cross-spectral matrix, Eqn. (10), used to form the correction to the multitaper PAC for a non-sinusoidal modulating oscillation. Thus, in this context, it is useful to use the Welch method to reduce the number of pulses per section to close to one. This can be accomplished in many cases based upon the expected rate of occurrence of pulses. One example is slow-waves during stage two non-rapid eye movement sleep; they occur at a rate of approximately 1-2 per minute [78].

## 6 Assessment of Statistical Significance

Let *y* be a zero-mean random process with a lagged-covariance independent of the absolute time, *t*; that is, the autocovariance sequence, *r*, is independent of *t*:

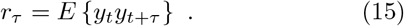

Under mild conditions, the eigencoefficient, Eqn. (2), is asymptotically circularly-symmetric normal; see for example, [58, 73, 79, 80]. Then the probability of |*Ĉ*(^*pac*^)(*f*)|^2^ exceeding the observed square-magnitude multitaper PAC,|*Č*^(*pac*)^(*f*)|^2^, conditioned upon the population coherence *C* equaling zero is,^4^

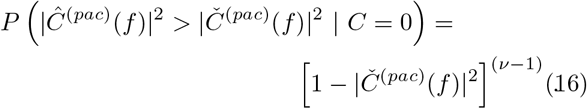

Here, Eqn. (16) is valid for frequencies excluding DC and Nyquist frequency, i.e. *f* ≠ 0,1*/*2Δ Hz, and *v* is equal to the degrees of freedom. This latter quantity is equal to the number of tapers times the number of trials. Eqn. (16) is equal to the p-value of a one-sided hypothesis test between the null hypothesis of no phase-amplitude coupling and the alternate hypothesis *C* > 0. Transformed p-values computed using Eqn. (16) are shown on the right-hand-side of Fig. (13) for intracranial EEG recordings exhibiting slow-wave spindling PAC.

## 7 Simulation

### 7.1 Multitaper Instantaneous Amplitude: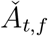

The estimate, 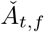, is tested on two signals, the first, an amplitude modulated sinusoid, and the second, a 5 ms pulse. Gaussian noise is added to both. The result is shown in Fig. (4) where the 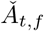, this time with *w* set equal to 1 in Eqn. (29), follows the amplitude of the modulated oscillation, and is nearly identical to the estimate computed using the Hilbert method. Both estimates depicted in Fig. (4) are also large for the broadband 5 ms pulse, which is undesirable. The situation can be mitigated, as shown in Fig. (5). In this case, the weight *w* is as specified in Eqn. (37), resulting in a nearly zero instantaneous amplitude when tested on the 5 ms pulse. This robust estimate is biased slightly low relative to the Hilbert instantaneous estimate when the signal is the modulated oscillation.

**Figure 4:**
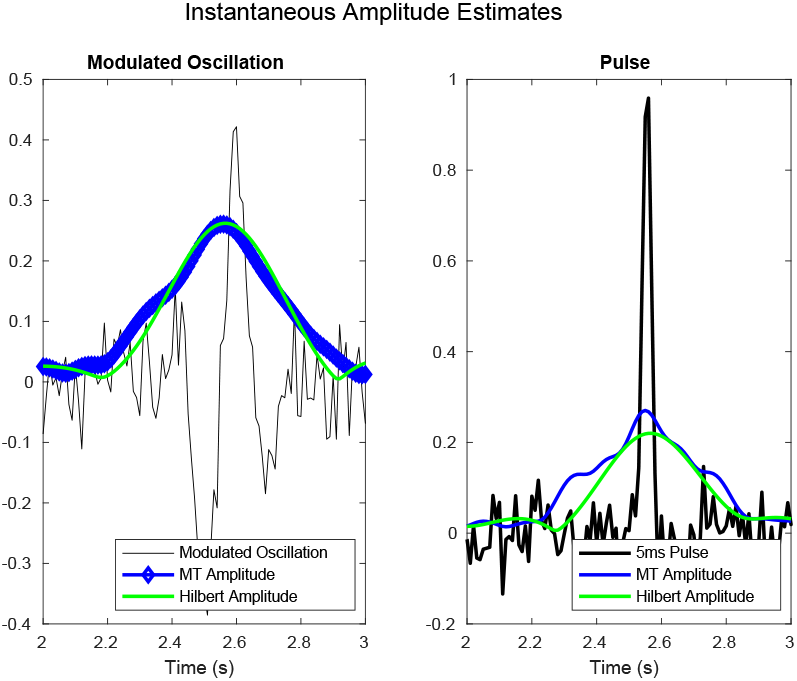
The unweighted multitaper instantaneous amplitude (*w* = 1) correctly estimate the amplitude of the oscillation (left), as does the estimate computed with the Hilbert transform. Both the multitaper and the Hilbert estimates respond to the 5 ms pulse (right). The units associated with the vertical axes of both plots (left and right) are arbitrary but equal.

**Figure 5:**
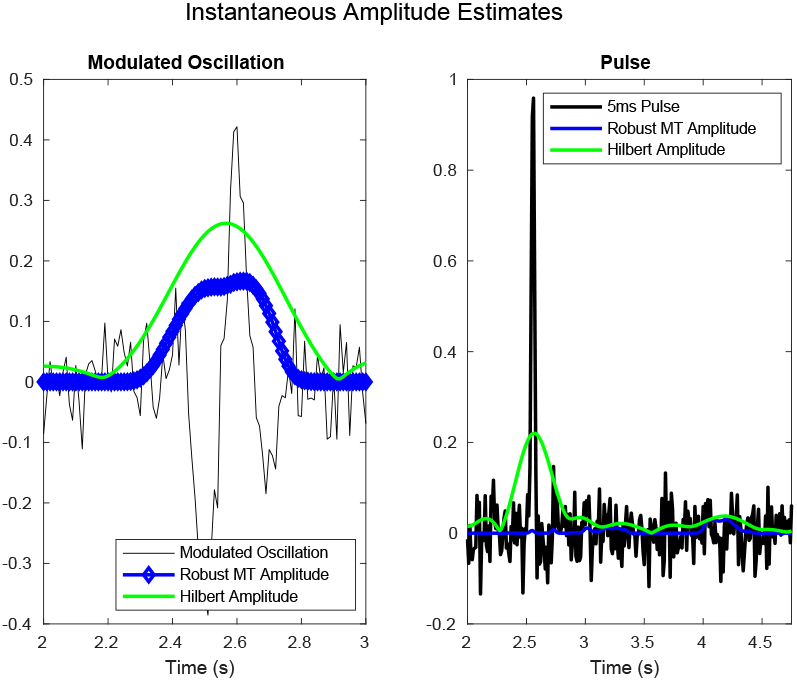
The weighted multitaper instantaneous amplitude estimate (blue curve, *w* specified as in Eqn. (37)) is biased low (left), but does not respond to the nonsinsoidal high-frequency pulse signal (right), while the Hilbert estimate (green) behaves as in Fig. (4), and responds to both signals. The units associated with the vertical axes of both plots (left and right) are arbitrary but equal.

### 7.2 Multitaper Phase Amplitude Coupling: 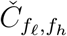

To test the accuracy of the proposed multitaper estimate in a challenging, yet realistic scenario, 100 experiments’ worth of data are created. For each simulated experiment, two modulating oscillations, oscillating at 2.2 Hz, and 6 Hz, respectively, are multiplied by Gaussian envelopes so that each modulating pulse persists for a few cycles when present, and occur at a rate of 1 every ten seconds, or 3 per each of the 30 second trials.^5^ Each experiment is comprised of 20, 30 second trials. One-over-f noise is generated following the filtering method presented in [81, §2.2.7], scaled to have a standard-deviation equal to about 1/4 of the maximum amplitude of the modulating pulses. Finally, modulated pulses are created and added to the synthetic recordings. The first modulated signal is generated by bandpass filtering white-noise to the interval (20 Hz, 30 Hz), and multiplying the filter output by a Gaussian with four standard deviations equal to 250 ms to generate a 2 Hz bandwidth pulse. The second modulated signal is created by bandpass filtering white noise to the interval (40 Hz, 60 Hz), and multiplying by a Gaussian with four standard deviations equal to 100 ms to generate a 5 Hz bandwidth pulse. These modulated signals, one for each occurring modulating pulse, are added to the synthetic data. Figure (6) shows the simulated data for experiment one. From the data shown in Fig. (6), the multitaper estimate of PAC, *C*^(*pac*)^, is computed, along with the PAC estimate computed using the Hilbert transform. The estimates are shown in Fig. (7) for modulating frequencies ranging from zero to ten Hertz, and for, modulated frequencies ranging from ten to seventy Hertz. The proposed multitaper estimate exhibits greater response, specificity, and accuracy than the PAC estimate computed using the Hilbert transform.

**Figure 6:**
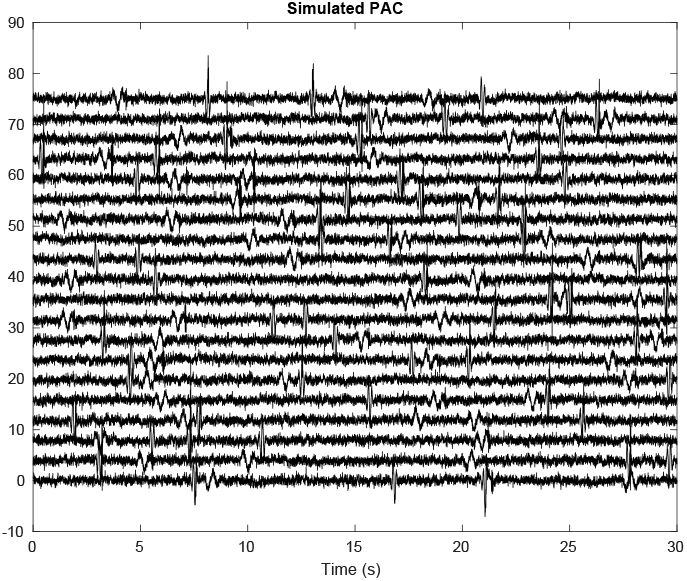
Simulated data exhibiting phase-amplitude coupling. Temporally-intermittent oscillatory pulses are apparent. Associated with the oscillatory pulses oscillating with a frequency of 2 and 6 Hz are oscillatory pulses oscillating with 25 and 50 Hz frequencies; respectively. These latter two oscillatory pulses are not phase-locked to the 2 and 6 Hz pulses; however, their amplitude is consistently timed to the lower frequency pulses.

**Figure 7:**
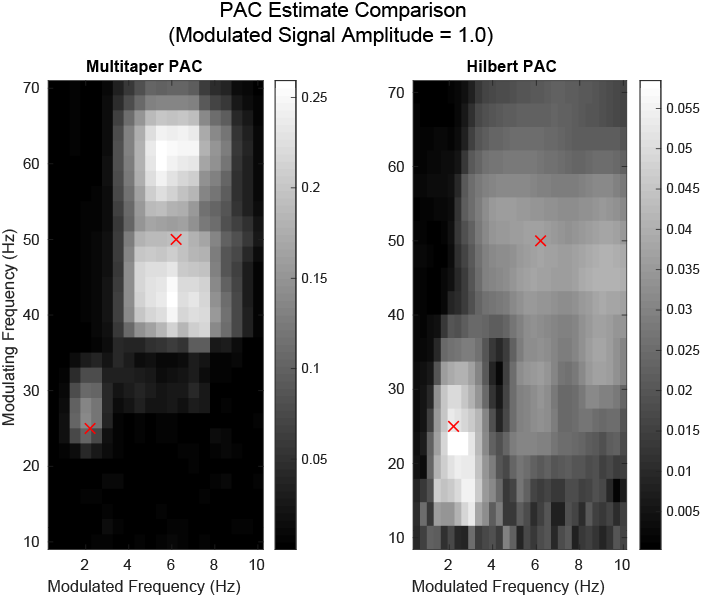
Left: Multitaper phase-amplitude coupling (PAC) estimate |*C*^(*pac*)^|. Right: PAC estimate computed using the Hilbert transform. The estimates are computed from the data depicted in Fig. (6). The proposed multitaper estimate exhibits greater response, specificity, and accuracy than the PAC estimate computed using the Hilbert transform. The red crosses indicate the center of the modulating and modulated frequency intervals.

### 7.3 Resistance to Non-Sinusoidal Oscillation

To investigate the performance of the eigencoefficient correction (see Section (5)) in the computation of the multitaper phase-amplitude coupling estimate, Eqn. (5), a non-sinusoidal oscillation, *y*, is convolved with a binary pulse train with one’s separated by four periods of the non-sinusoidal oscillation *y*, and added to Gaussian white noise to obtain one of eighty simulated trials. Here, the non-sinusoidal pulse, *y*, indexed at time-index, *t*, is equal to,

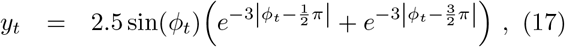

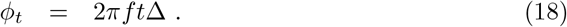

This results in the black curve in Fig. (8), which shows the first of the eighty trials. From the simulated data, the proposed eigencoefficient correction specified in Section (5) is computed, and the corrected eigencoefficients are used to reconstruct the time-series, yielding the blue curve also shown in Fig. (8). The correction reduces the large excursions from a sinusoidal oscillation exhibited at the extrema of the sawtooth waveform. From this simulated sawtooth data which, by construction, exhibits no phase-amplitude cross-frequency coupling beyond that which can be explained by phase-phase coupling, two multitaper PAC estimates are computed and shown in Fig.(9). The first estimate is the proposed multitaper PAC estimate (Eqn. (5)), and the second is the corrected estimate computed using Eqn. (9). To demonstrate the ability of the non-sinusoidal corrected multitaper estimate to detect non-spurious PAC associated with a non-sinusoidal modulation, this first simulation is repeated, but in this case, additional PAC is added by adding white noise filtered to the frequency interval (21 Hz, 26 Hz), and multiplied with a Gaussian pulse train; each pulse having a standard deviation equal to 1*/*4*f* seconds, and spaced by 4*/f* seconds (the same periodicity as the sawtooth oscillatory pulse train). The first trial of the resulting data (black) and the multitaper corrected reconstruction (blue) are shown in Fig. (10). Again, multitaper phaseamplitude coupling estimates are computed (Fig. (11)). The corrected multitaper PAC estimate is not close to zero for all frequencies plotted, but instead, reveals the added phase-amplitude coupled component present in the simulated data (shown in Fig. (10)).

**Figure 8:**
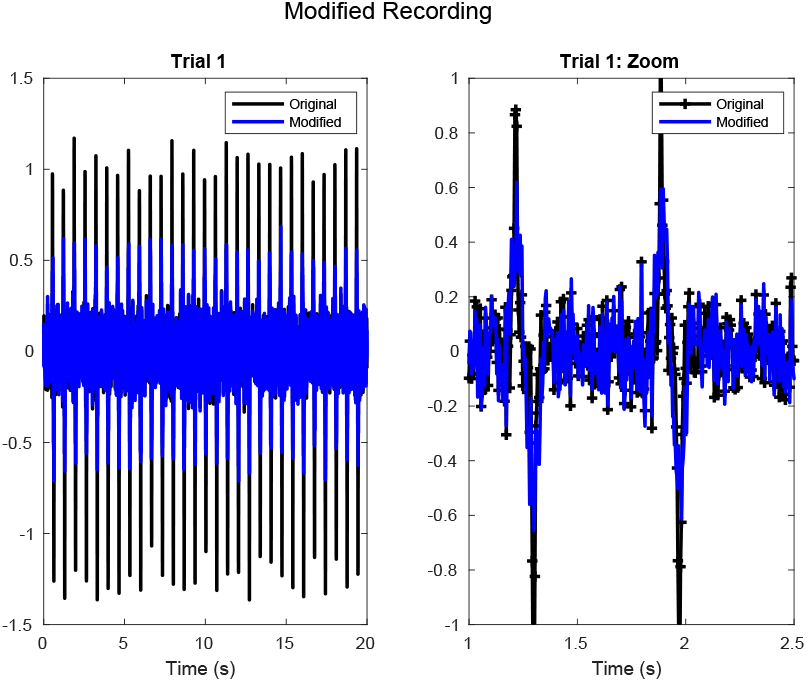
One trial of the simulated sawtooth data (black) and its multitaper reconstruction after removing estimated PPC (blue). No amplitude modulated components are added to the periodic, non-sinusoidal, sawtooth oscillations (black). The multitaper reconstruction (blue), computed using the corrected eigencoefficients (see Section (5)), reduces the absolute extremes of the sawtooth oscillation.

**Figure 9:**
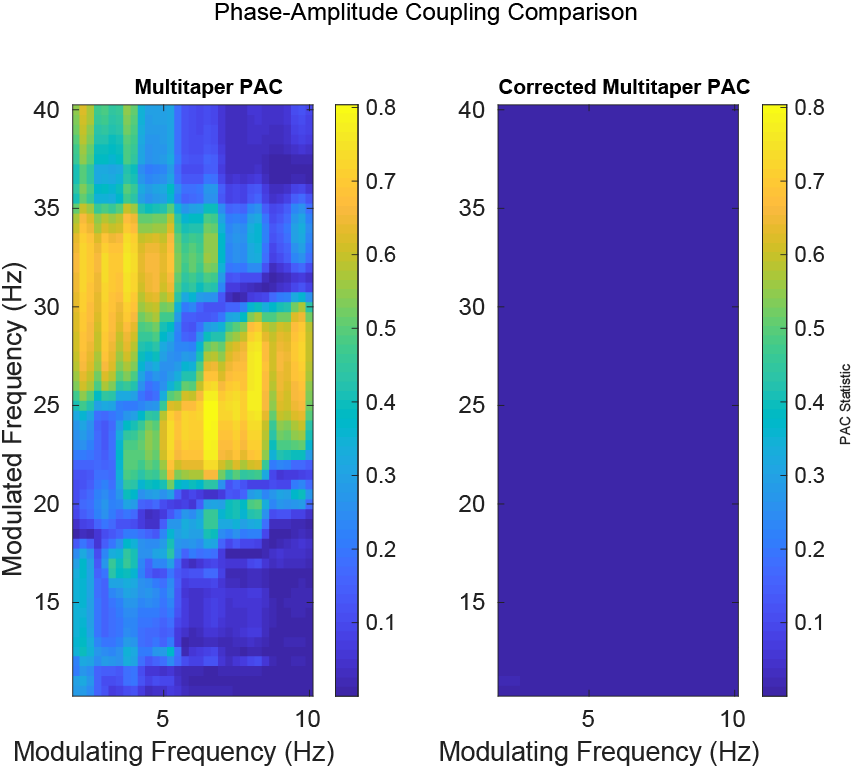
The proposed correction for non-sinusoidal modulation removes spurious PAC. The proposed multitaper phase-amplitude coupling estimate, Eqn. (5), responds to the non-sinusoidal phase-amplitude cross-frequency coupling inherent in non-sinusoidal oscillations (left). The proposed correction, see Section (5), greatly reduces the undesirable response (right). The corrected PAC estimate is not zero; however, this cannot be seen on the scale displayed.

**Figure 10:**
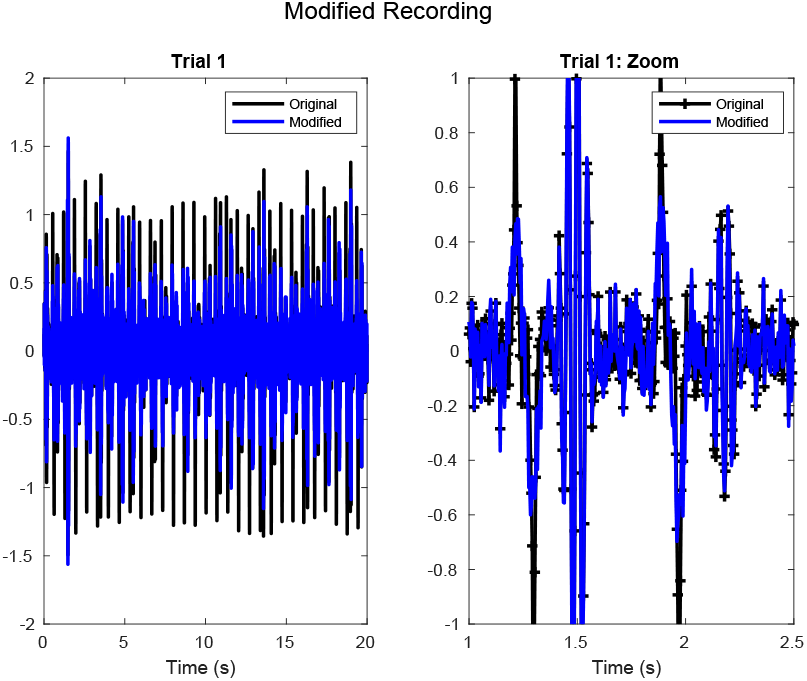
One trial of the simulated sawtooth data (black) and its multitaper reconstruction after removing estimated PPC (blue). Amplitude modulated components have been added to the periodic, non-sinusoidal, sawtooth oscillations (black), one for each saw-tooth oscillation. The amplitude modulated components are Gaussian-enveloped, white noise filtered to have signal energy within the frequency interval (21 Hz, 26 Hz). Thus, the phase of the resulting modulated noise is, ignoring negligible temporal correlation, random from pulse to pulse, while the envelope is time-locked to the non-sinusoidal sawtooth oscillations.

**Figure 11:**
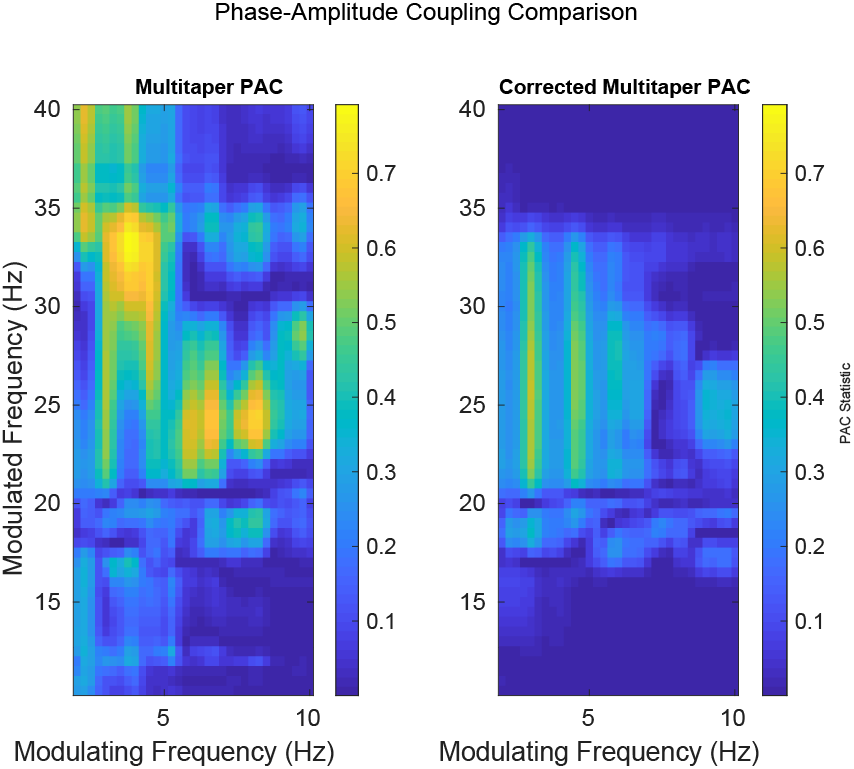
The proposed correction for non-sinusoidal modulation reveals PAC obscured by non-sinusoidal modulation. The proposed multitaper phase-amplitude coupling estimate (Eqn. (5)), responds to the non-sinusoidal phase-amplitude cross-frequency coupling inherent in non-sinusoidal oscillations, as well as to non-spurious PAC (left). The proposed correction (see Section (5)), greatly reduces the undesirable response, and reveals PAC not-attributable to the sawtooth oscillation alone (right). Compare with Fig. (9) for multitaper PAC estimates computed from sawtooth oscillations without an extra phase-ampltude coupled component present.

## 8 Intracranial EEG

Intracranial EEG recordings from Carilion Roanoke Memorial Hospital, Virginia were obtained from an epileptic patient during non-REM sleep. Bipolar referencing (aka, ‘subtraction’) between two contacts in the middle frontal gyrus were used to compute the introduced multitaper phase-amplitude cross-frequency coupling (PAC) estimate using Eqn. (5). ^67^ The estimate is shown in Fig. (12). For comparison, the PAC estimate computed using the Hilbert transform is also shown (Fig. 12, left). The modulated oscillation bandpass filter passband is specified to be equal to (*f*_*h*_ − 1.1*f*_*l*_, *f*_*h*_ + 1.1*f*_*l*_); consistent with the filter passband recommended in [42]. The proposed multitaper PAC estimator is less smeared in frequency than the Hilbert estimate.

**Figure 12:**
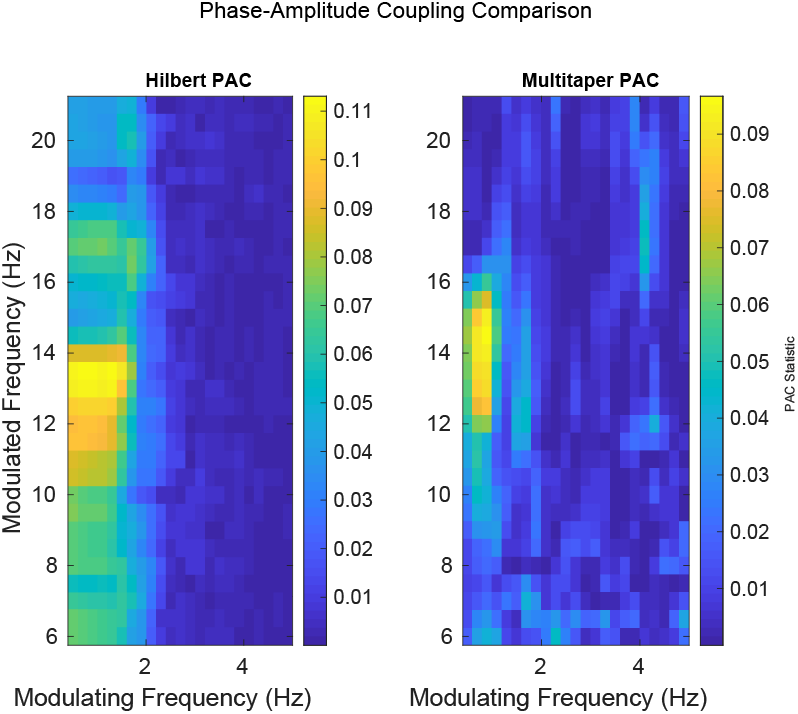
Comparison between the Hilbert PAC statistic and the proposed multitaper PAC statistic. The multitaper estimate exhibits greater specificity and less smearing in frequency. The peak of the multi-taper PAC estimate occurs at a modulated frequeny that is around 2 Hz greater than the associated Hilber PAC estimate.

## 9 Discussion

Two nonparametric, multitaper estimates of phase-amplitude coupling are introduced. The first of these estimates is susceptible to non-oscillatory modulating oscillations (Eqn. (5)), while the second estimate is resistant (§5). Both estimates are resistant to broadband pulses, owing to the introduction of a pulse-resistant multitaper estimate of the instantaneous amplitude. An expression for theoretical p-values is given in Section (6). The estimates are tested in simulation (§7) and on intracranial EEG data (§8), where the stated resistances are demonstrated. The multitaper PAC estimates are shown to be superior to the comparable PAC estimate calculated using the Hilbert transform in these cases. The bandwidth of the proposed PAC estimates is set first by the number of constant periods of the modulated, or high-frequency oscillation (3 periods of oscillation in this work), and for the multitaper coherence between the low and high frequencies, the bandwidth parameter *W* is left to be discovered by exploratory analysis. In this work, this value is set to 0.5 Hz.

This work addresses major concerns when estimating phase-amplitude coupling. The first two concern spurious PAC due to (i) non-oscillatory modulation, and (ii) to contaminating broadband activity, such as an interictal event. Further, by introducing these multitaper PAC estimates, multitaper variance reduction is achieved, as is the availability of an asymptotically valid p-value for testing for PAC significance. This latter quantity does depend upon the weak-sense stationary assumption; however, as can be seen in Fig. (13) it is effective on real intracranial data. Finally, weak-sense stationary models of phase-amplitude coupling are reported in [81].

**Figure 13:**
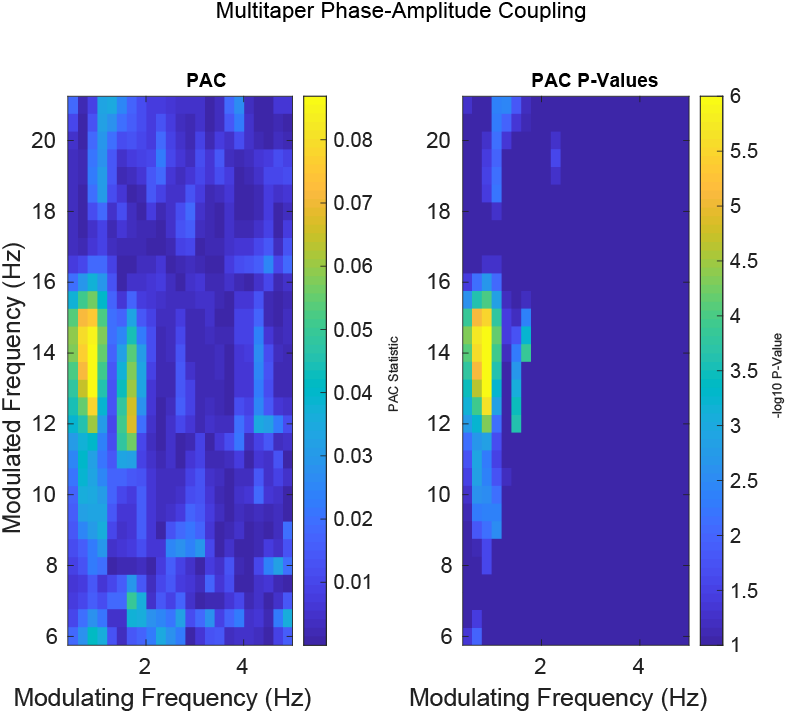
The phase-phase cross-frequency correction for nonsinusoidal phase-amplitude coupling (see §5) reveals extra structure in intracranial data. Left: The phase-phase cross-frequency coupling corrected multitaper estimate. Compare with the uncorrected estimate shown in the right-hand image in Fig. (12). Right: The −log_10_ transformed p-values computed using Eqn. (16). These p-values have not been corrected for multiple comparisons. The p-values account for random statistical fluctuations and suppress spurious activity.

## Acknowledgements

SV acknowledges support from the U.S. Army Research Office under award number ARO W911NF-17-1-0300.

Research reported in this publication was also supported in part by the National Center for Advancing Translational Sciences of the National Institutes of Health under Award Number UL1TR003015. The content is solely the responsibility of the authors and does not necessarily represent the official views of the National Institutes of Health. We thank Anna E. Vijayan for carefully reading and editing the manuscript.

## A Instantaneous Amplitude, 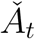

The estimate, 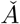 is derived by first considering the timedependent eigencoefficient,

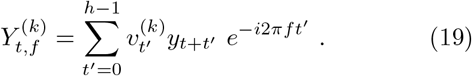

Let

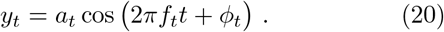

Then, substituting Eqn. 20 into Eqn. 19 results in,

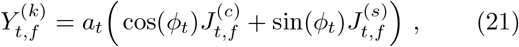

where

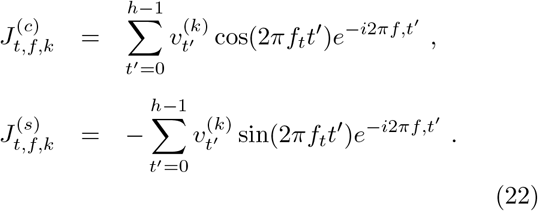

For 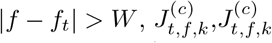 are approximately equal to zero. For *f* equal to *f*_*t*_:

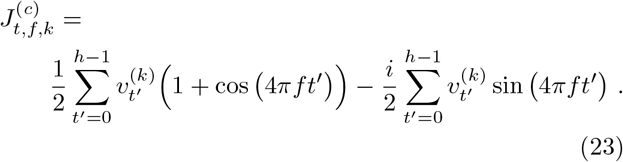

For |*f*| < *W*,

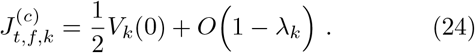

Here *λ*_*k*_ is the eigenvalue associated with the *k*^*th*^ DPSS, and *V*_*k*_(*f*) is the discrete-time Fourier transform of *v*^(*k*)^ evaluated at frequency *f*. I.e.:

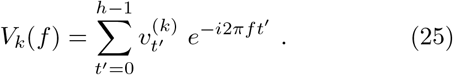

Continuing, for *f* equal to *f*_*t*_,

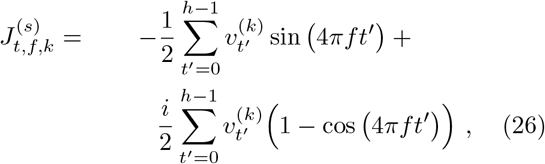

and for |*f* | > *W*

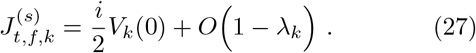

Substituting Eqns. (24), (27) into Eqn. (19), and evaluating with *f* = *f*_*t*_, |*f* | > *W*, results in the time-dependent eigencoefficient,

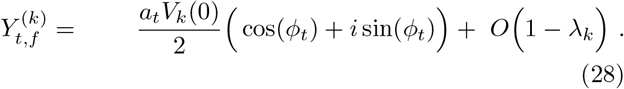

Eqn. (28) can be used to estimate both the instantaneous amplitude, *a*_*t*_ and the instantaneous phase, *ϕ*_*t*_. Let the *k*^th^ entry of **x** equal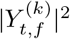. The proposed multitaper estimate, 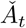 of *a*_*t*_ is equal to,

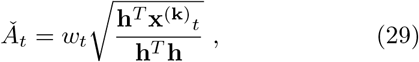

where the *k*^th^ element of **h** is equal to 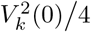 and the weight, *w*, is specified in the next appendix; see Eqn. (37) A multitaper estimate of the instantaneous phase, *ϕ*_*t*_, is

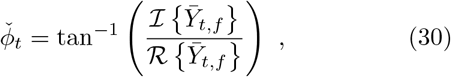

with 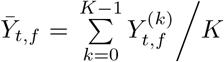 The weight, *w*, in Eqn. (29), is used to bias the instantaneous amplitude estimate to zero in the case of narrow-band, yet non-sinusoidal signal energy (see B).

## B F-test for Sinusoidal Oscillation

Following [58, 76, 82], an F-statistic relating the signal-power explained by a sinusoid within the linear, normed, vector space spanned by the in-band energy concentrated DPSS sequences modulated to a test frequency *f*, to the signal-power that is unexplained in this vector-space. This ratio is suppressed, for example, when two sinusoids oscillating at frequencies within the frequency interval (*f* −*W, f* + *W*) are present, and is large, when the narrowband signal energy is largely due to that of a sinusoid oscillating with a frequency belonging to the interval. Let the *t*^th^ row and *k*^th^ column of **C** equal the *k*^th^ modulated Slepian^8^. That is,

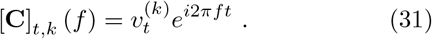

Let **1** equal the vector of 1’s. Modulating this vector to frequency *f* results in **1**_*f*_:

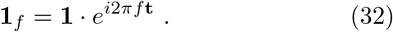

Here *·* is the Hadamard, or element-by-element vector product. Resolving **1**_*f*_ onto the column space of **C** results in,

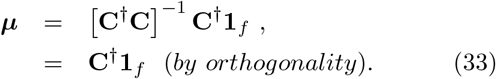

Similarly, modulate the observed recording, **y**:

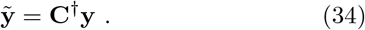

Note that 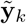 is equal to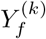, the *k*^th^ eigencoefficient (see Eqn. (2)). Now, the projection matrix onto the sinusoid in the column space of C is equal to,

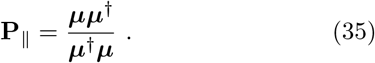

The F-statistic, *F*, is,

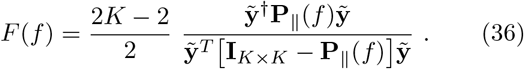

Under mild assumptions [58, 73, 79, 80], the Fourier coefficients of a recording are asymptotically circularly symmetric normal, and the ratio in Eqn. (36) is asymptotically F-distributed with 2 degrees of freedom in the numerator and 2*K* − 2 degrees of freedom in the denominator. P-values associated with this statistic are computed to obtain *p*_*f*_. The weight, *w*_*f*_ associated with *p*_*f*_, is set equal to,

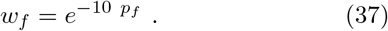

Thus, a p-value equal to 0.1 is suppressed one e-fold. This weight is used to down-weight non-sinusoidal contributions to the multitaper estimate of the instantaneous amplitude (see Eqn. (29)).

## Footnotes

1 The zero^th^ order DPSS is specified by the number of elements in the sequence, *n*, and by its bandwidth parameter, *W*. Of all sequences of length *n*, it is maximally energy-concentrated within the interval (−*W, W*). The *m*^th^ order DPSS is, of all sequences of length *n* that are orthogonal to the (*m* 1)^st^ order DPSS, the sequence that is maximally energy-concentrated within (−*W, W*) [56].

2 Data tapers reduce the influence of the ends of a time-series on the spectrum estimates. This is, roughly speaking, equivalent to discarding data and results in an increase in individual estimator variance. Averaging these estimates controls this variance increase.

3 There is some confusion regarding nomenclature in the literature. Here, we refer to | *Č*_*pac*_| as an estimate of the “coherence” between *f*_*£*_ and the instantaneous amplitude of *f*_*h*_; but, note the analogous terminology using “coherency”, and “complex coherency”, in, for example, the discussion surrounding Eqn. (9.1.35) in [73, p. 661].

4 Derived from Eqn. (2.6), p. 253 of [79].

5 A four modulating-pulse cycle refractory period is introduced by randomly removing pulses until only one pulse occurs in any given four cycle interval of time.

6 After decimation to a 250 Hz sample rate, and notch filtering to remove 60 Hz oscillations and harmonics.

7 Filtering is acausal to remove phase effects. Specifically, the MATLAB command, filtfilt() is employed for all filtering. In addition, edge effects are discarded from the filtered time-series prior to analysis.

8 Synonymous with the discrete prolate spheroidal sequence; the sequence is named after David Slepian, who studied them extensively[56].

